# DNMT3L interacts with Piwi and modulates the expression of piRNAs

**DOI:** 10.1101/2023.09.24.558918

**Authors:** Ramisetti Rajeev, Rakesh Mishra, Sanjeev Khosla

## Abstract

The epigenetic modulator, DNMT3L, has been shown to be involved in nuclear reprogramming. Previous work from our laboratory had shown the accumulation and inheritance of epimutations across multiple generations when DNMT3L was ectopically expressed in transgenic DNMT3L Drosophila. Here, we show interaction of DNMT3L with Piwi, a member of a family of proteins known to perform their function through their association with piRNAs. Importantly, DNMT3L expression caused a significant alteration in the piRNA profile across multiple generations in transgenic Drosophila. As piRNAs are known to be passed on from one generation to another, we believe that the DNMT3L through its interaction with Piwi act to modify the inherited pool of piRNAs, which in turn allows accumulation of aberrant epigenetic modifications in subsequent generation. Furthermore, we show the interaction of DNMT3L with Histone H1, a non-core histone involved in higher order chromatin organisation. In light of these observations, it is proposed that in addition to its role in modulating core histone modifications, DNMT3L allows for inheritance of non-genetic information through its collaboration with Piwi, piRNAs and histone H1.

## Introduction

The complex interaction between epigenetic writers, erasers, readers and epigenetic modifications such as DNA methylation, histone modifications and small non-coding RNA forms the basis of epigenetic inheritance [1, 2]. Several studies have revealed the inheritance of non-genetic information, including epigenetic modifications through gametes [3]. Usually, germ cells are known to undergo epigenetic reprogramming by erasure of existing and reestablishment of new epigenetic marks [4]. Any cues that influence this epigenetic reprogramming would ultimately affect or influence the zygote and the progeny in the next generation. If parental germ cells acquire aberrant epigenetic information due to changes in the environment such as stress, diet, infections, etc in the parents, these would be inherited by the progeny [5]. Thus, epigenetic inheritance is based on dynamic nature of epigenetic modifications and unlike genetic code, responds to both endogenous (developmental cues) and exogenous (environment) stimuli [6]. In evolutionary terms, it has been postulated that epigenetic gene regulation gives an advantage to organisms by adjusting to the continually varying environment and provides a vast range of phenotypes for the same stimuli [7, 8].

Studies on various model organisms have revealed that in addition to inheritance of DNA and histone modifications, small non-coding RNAs such as siRNAs, miRNAs, and piRNAs also form vectors for inheritance of acquired non-genetic information [9, 10]. piRNAs and Piwi proteins are associated with the protection of the genome in the germline from transposable elements and have been shown to be inherited and inducer of transgenerational epigenetic silencing[11, 12].

Dynamic addition and erasure of DNA methylation plays a crucial role during epigenetic reprogramming. DNMT3L is an epigenetic modulator, belonging to the *de novo* methyltransferase family that also includes DNMT3A and DNMT3B. DNMT3L lacks key amino acid residues in the catalytic domain and is unable to directly methylate DNA [13, 14]. However, through its direct interaction with DNMT3A, DNMT3B and histone H3, DNMT3L participates in the recruitment of DNA methylation machinery in response to histone modifications [15, 16]. DNMT3L is also a key protein associated with germ cell reprogramming as knockout of Dnmt3l in mice has been shown to cause a broad range of developmental abnormalities and genomic imprinting defects [17].

Previous work from our laboratory uncovered the role of DNMT3L in nuclear reprogramming upon its ectopic expression in transgenic Drosophila [18]. The reprogramming was gradual and progressive and only in the 5^th^ generation transgenic Drosophila larvae were the melanotic tumors observed. This was concomitant with progressive accumulation of epimutations, in the form of aberrant H3K4me2, H3K4me3 and H3K36me3 histone modifications. Progeny of crosses between the Piwi mutant (piwi^06843cn1/CyO^; ry^506^) and DNMT3L did not show any melanotic tumors suggesting a role for the Piwi pathway in the inheritance of epimutations in DNMT3L expressing Drosophila.

Present study explores the role of Piwi protein and piRNAs in DNMT3L mediated epigenetic inheritance by examining the association of DNMT3L with histones and small RNA, especially piRNA. We show that the interaction of DNMT3L with histone H1, H3 and Piwi protein leads to change in the profile of several groups of small RNAs (sRNAs) including piRNAs in transgenic DNMT3L Drosophila. Our study provides for an important mechanism for integration of DNA methylation, histone modifications, higher order chromatin organisation and small RNA cues into the inheritance of non-genetic information across generations.

## Results

### DNMT3L interacts with Piwi protein

Previous study from our laboratory had shown that ectopic expression of DNMT3L causes a progressive misregulation of multiple genes concomitant with a gradual reduction of specific active histone marks. Within the subset of misregulated genes, it was observed that all the Piwi group of proteins were upregulated. Piwi gene itself was highly upregulated in the 5^th^ generation. Moreover, progeny from crosses set up between 5^th^ generation DNMT3L expressing flies and Piwi mutant flies did not show any tumour phenotype, suggesting genetic interaction with DNMT3L [18]. Piwi and its associated RNAs (piRNA) are known to be involved in the inheritance of epigenetic information across generations in Drosophila [19, 20]. Studies from other laboratories have also indicated that DNMT3L and Piwi protein works in the same pathway for targeting of small RNAs to specific loci in germ cells [21, 22]. Therefore, we were keen to explore whether Piwi and DNMT3L physically interact with each other. Recombinant 6X His-Flag-DNMT3L was expressed and purified from *E coli* BL21 strain and incubated with adult Drosophila cell lysate followed by affinity pull-down using Talon beads. Piwi protein was found to be enriched in DNMT3L pull-down fraction (Supplementary Figure S1). Piwi protein is normally expressed at higher level only in the germ cells. To further validate DNMT3L and Piwi interaction, DNMT3L immunoprecipitation was performed on ovary lysate from DNMT3L transgenic flies using DNMT3L antibody. The DNMT3L-bound proteins were western blotted and probed with anti-Piwi antibody. Piwi protein was detected in the DNMT3L immunoprecipitated fraction (Figure 1A). Vice versa, immunoprecipitation was performed with Piwi antibody and western blot for the bound proteins was probed for DNMT3L enrichment (Figure 1B). Both these experiments revealed that DNMT3L and Piwi can interact.

**Figure 1:**
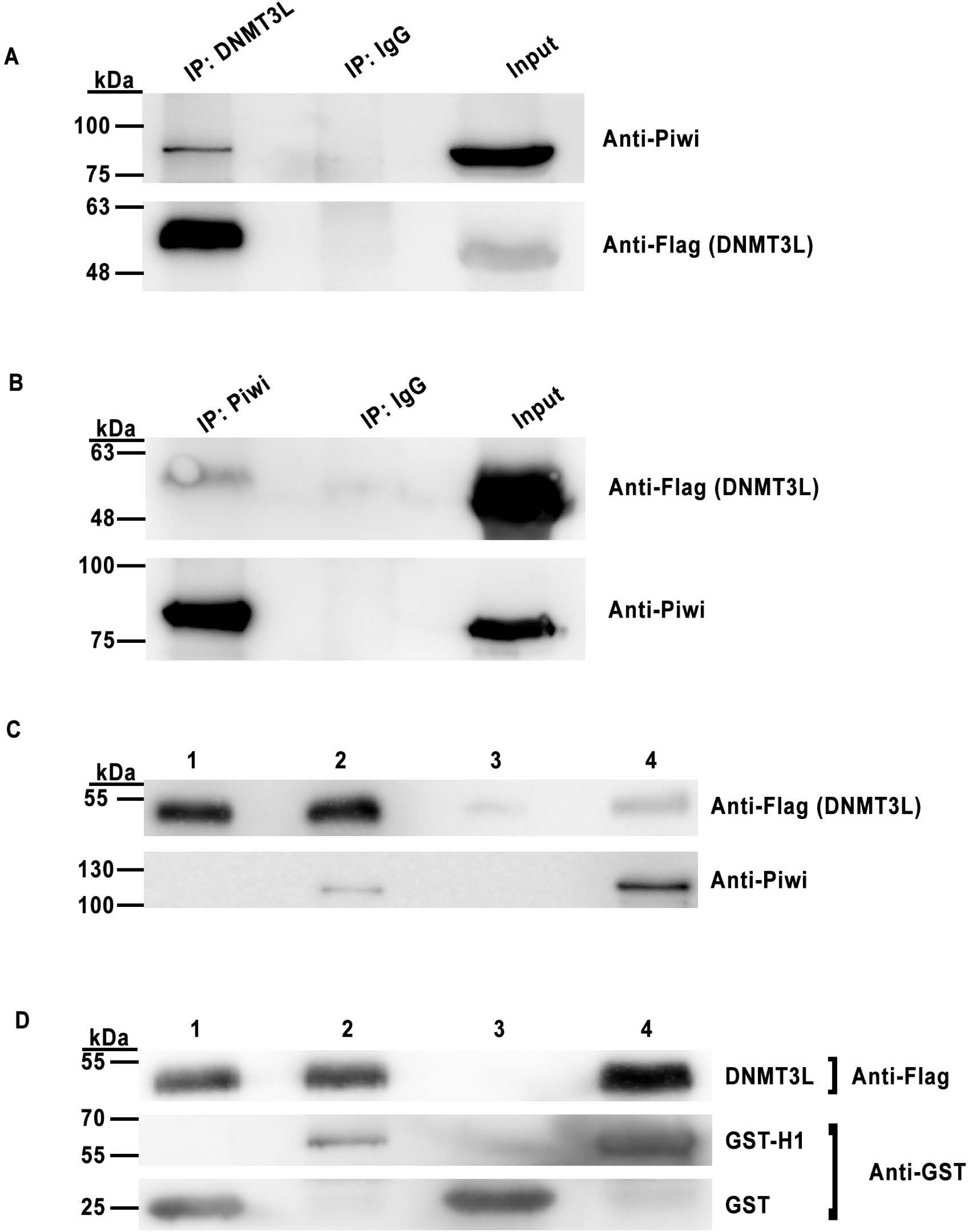
DNMT3L protein interacts with Piwi and histone H1. A. Western blot analysis was performed for DNMT3L-associated proteins in the transgenic FLAG-DNMT3L Drosophila ovary lysate subjected to immunoprecipitation (IP) using anti-DNMT3L. The blot was probed with anti-Piwi antibody. IP with anti-rabbit IgG was used as control. B. Same as (A.) but IP with anti-Piwi antibody and probing with anti-Flag antibody. C. To demonstrate direct DNMT3L and PIWI interaction, pulldown was performed with *E.coli* purified recombinant GST-Piwi and 6X His-Flag-DNMT3L. As a control 6X His-Flag-DNMT3L interaction was examined with GST alone. Western blot analysis with respective antibodies. Lane 1-Input (GST + 6X His-Flag-DNMT3L), 2-Input (GST Histone H1 + 6X His-Flag-DNMT3L), 3-Pulldown with GST for GST+6X His-Flag-DNMT3L and 4. Pulldown with GST for GST-Piwi+6X His-Flag-DNMT3L. DNMT3L and PIWI specific bands were detected at their respective size. D. DNMT3L protein interacts with Histone H1. To demonstrate direct DNMT3L and histone H1 interaction, pulldown was performed with *E.coli* purified recombinant 6X His-Flag-DNMT3L and GST-Histone H1 or control GST alone. Western blot analysis was performed with either FLAG (DNMT3L) or GST (histone H1). Lane 1-Input (GST + 6X His-Flag-DNMT3L), 2-Input (GST-Histone H1 + 6X His-Flag-DNMT3L), 3 – Pulldown for GST on GST+6XHis-Flag-DNMT3L and 4-Pulldown for GST-Histone H1 on GST-Histone H1+ 6X His-Flag-DNMT3L.

To test whether this interaction was direct or indirect, pulldown with bacterial purified proteins was performed. recombinant GST-Piwi and 6X His-Flag-DNMT3L proteins were separately purified from *E. coli* and co-incubated. GST-Piwi protein was pulled down using Glutathione agarose beads and the bound proteins were western blotted and probed with anti-Flag antibody to examine the enrichment of 6X His-Flag-DNMT3L. As shown in Figure 1C, DNMT3L was detected in the GST-PIWI pulldown fraction but not in pulldown with purified GST alone incubated with 6X His-tagged Flag-DNMT3L.

### DNMT3L interacts with Piwi-interacting protein Histone H1

Piwi protein interacts with histone H1 in a complex that regulates chromatin accessibility, especially to repress transposons [23]. Interestingly, in a mass-spectrometric analysis performed in our laboratory, to identify DNMT3L interacting proteins, histone H1 was identified [24]. As the Piwi-histone H1 containing complex is involved in the piRNA mediated transcriptional and post-transcriptional regulation, we sought to examine whether DNMT3L could also bind to histone H1. Recombinant 6X His-Flag-DNMT3L and GST-Histone H1, individually expressed and purified from *E. coli* were co-incubated. GST-Histone H1 was pulled down using Glutathione beads, the bound fraction was western blotted and probed with anti-Flag antibody to examine the enrichment of Flag-tagged DNMT3L. As shown in Figure 1D, DNMT3L was detected in the GST-Histone H1 pulldown fraction. Purified GST-alone incubated with 6XHis-Flag-DNMT3L was used as a control for the experiment.

### Expression analysis of small RNAs in transgenic DNMT3L Drosophila

We had previously hypothesized that the observation of progressive nuclear reprogramming leading to melanotic tumors in the 5^th^ generation of transgenic DNMT3L Drosophila was due to inheritance of accumulated epimutations over several generations [18]. As shown above, DNMT3L interacts with Piwi and histone H1, the known regulators of piRNA mediated epigenetic inheritance. Moreover, in a recent study, mouse Piwi protein PIWIL4 has also been shown to be involved in the demethylation of H3K4me2 [25]. In DNMT3L expressing transgenic Drosophila, the levels of active histone marks, H3K4me2, H3K4me3 and H3K36me3, had progressively decreased [18]. To investigate if any correlation exists between small RNA species and inheritance of epigenetic modifications in DNMT3L transgenic Drosophila, we performed a sRNA-seq (small RNA-sequencing) study to compare the profile of sRNAs especially piRNA in embryos and ovaries of transgenic DNMT3L Drosophila. sRNA was isolated from ovarian tissue and early stage two embryos of 1^st^ (G1) and 5^th^ (G5) generation Tubulin-Gal4-DNMT3L (DNMT3L expressing). sRNA from UAS-DNMT3L (control transgenic flies without DNMT3L expression) flies was used for the comparison. The obtained small RNA sequence reads (NCBI-PRJNA993000) were analysed using Unitas software [26] to find out the overall composition of small RNAs in each tissue.

The composition of small RNAs in the ovaries of 1^st^ and 5^th^ generation of DNMT3L expressing transgenic Tubulin-Gal4-DNMT3L Drosophila was different from control UAS-DNMT3L flies. The proportion of ovarian piRNAs from G1 flies was 52.1%, in G5 flies it was 49.5% whereas it was 47.3% in the control UAS-DNMT3L flies (Figure 2 & Table 1). The miRNA proportion (19.3%) was similar in control UAS-DNMT3L and G1 Tubulin-Gal4-DNMT3L but in G5 Tubulin-Gal4-DNMT3L flies the proportion was slightly higher at 20.8% (figure 2 & Table 1).

**Figure 2:**
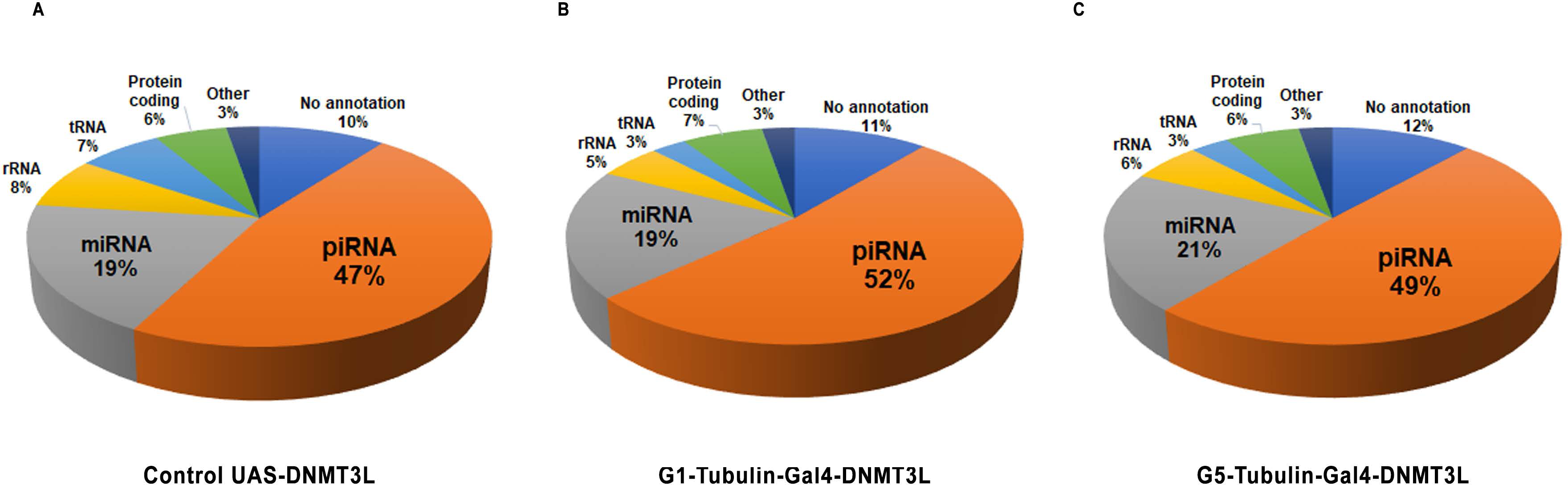
Composition of small RNAs (sRNA) in transgenic Drosophila ovary. **A.** Pie charts representing the percent distribution of overall composition of small RNAs (sRNA) in the ovary tissues isolated from **A**. UAS-DNMT3L control (without DNMT3L expression), **B**. 1^st^ generation and **C.** 5^th^ generation of Tubulin-Gal4-DNMT3L (DNMT3L expressing) transgenic flies as identified in Unitas software.

**Table 1:** Composition of small RNAs (sRNA) in transgenic Drosophila ovary.

| Small RNA type | Control Uas-DNMT3L | G1-Tubulin-Gal4-DNMT3L | G5-Tubulin-Gal4-DNMT3L |
| --- | --- | --- | --- |
| piRNA | 47% | 52% | 49% |
| miRNA | 19% | 19% | 21% |
| rRNA | 8% | 5% | 6% |
| tRNA | 7% | 3% | 3% |
| Protein coding | 6% | 7% | 6% |
| Other | 3% | 3% | 3% |
| No annotation | 10% | 11% | 12% |

The proportion of piRNAs in Drosophila embryos was substantially lower than ovaries. In embryos, piRNAs proportion in G5 Tubulin-Gal4-DNMT3L was significantly higher (25.1%) as compared to the control UAS-DNMT3L flies (13.7%, Supplementary Figure S2). The proportion of miRNA also decreased from 21.4% to 17.4% in G5 Tubulin-Gal4-DNMT3L embryos as compared to control (Supplementary Figure S2).

In the piRNAs detected in the ovarian tissue, approximately 17% (8% overexpressed and 9% downregulated) were differentially expressed in the ovaries of G5 Tubulin-Gal4-DNMT3L transgenic flies as compared to UAS-DNMT3L flies. On the other hand, approximately 48% (23.6% overexpressed and 24.5% were downregulated) were differentially expressed in G1 Tubulin-Gal4-DNMT3L transgenic flies as compared to UAS-DNMT3L flies (Table 2). A comparison of G1 and G5 Tubulin-Gal4-DNMT3L transgenic flies showed that approximately 9% piRNAs were upregulated and 17% were down regulated in G5 flies (Table 2). Hierarchical clustering of small_RNA-seq samples to examine differences in piRNA expression showed a clear separation of expression level of piRNAs in control, G1 and G5 Tubulin-Gal4-DNMT3L transgenic flies (Figure 3A). The length distribution of small RNAs was also found to be affected (Supplementary Figure S3).

**Figure 3:**
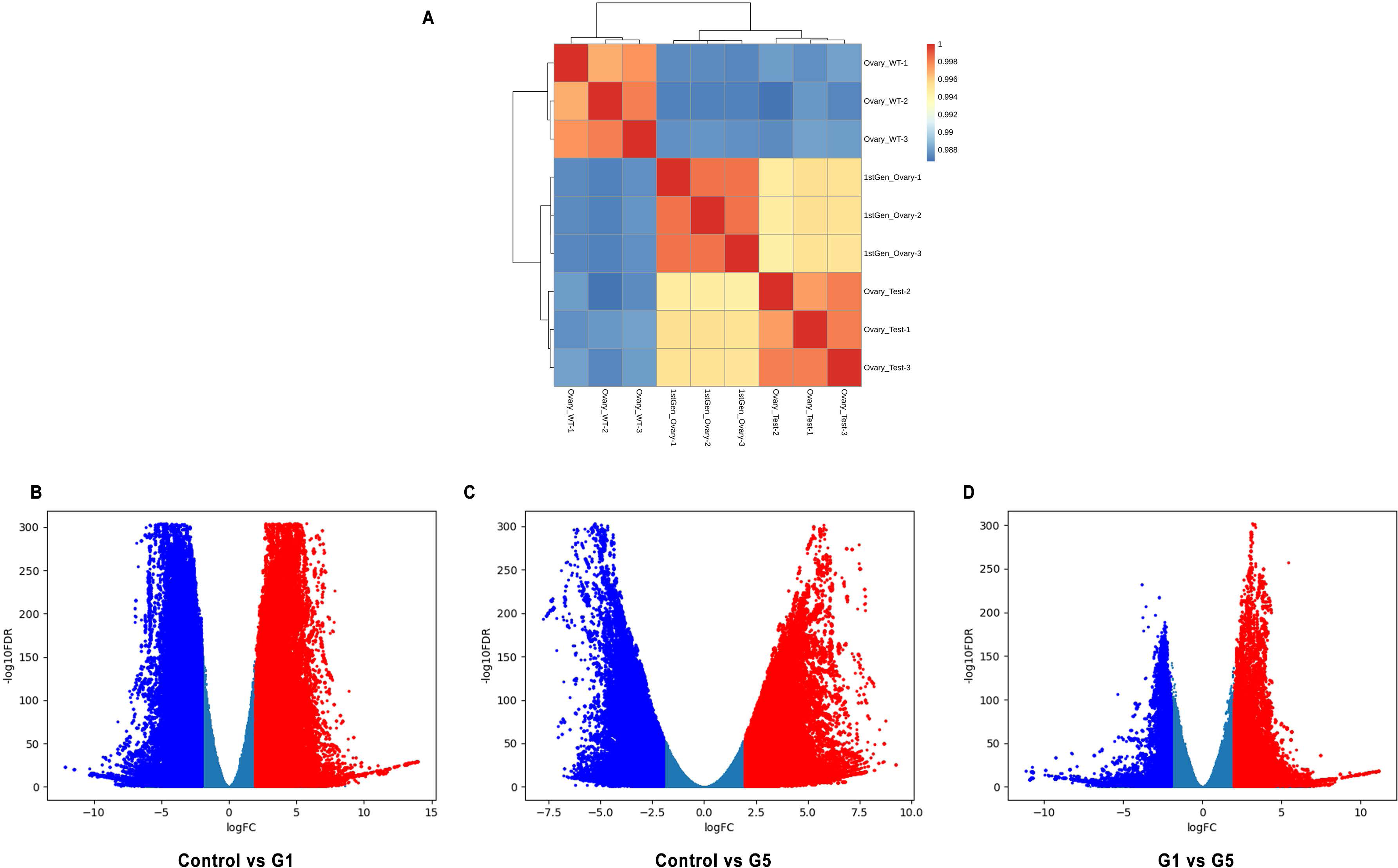
Comparison of differentially expressed piRNAs. A.Hierarchical clustering plot was generated to define correlation of differential piRNA expression derived from ovarian tissue of control UAS-DNMT3L and Tubulin-Gal4-DNMT3L transgenic flies from G1 and G5 generation. WT – control UAS-DNMT3L, Ovary-Test G5 Tubulin-Gal4-DNMT3L flies, and 1st gen-ovary - refer to G1 Tubulin-Gal4-DNMT3L flies. 1, 2 and 3 for each sample refers to the three biological replicates. Volcano plot comparison of piRNA profile between: B. control UAS-DNMT3L and G1-Tubulin-Gal4-DNMT3L; C. control UAS-DNMT3L and G5-Tubulin-Gal4-DNMT3L; D. G1-Tubulin-Gal4-DNMT3L and G5-Tubulin-Gal4-DNMT3L. Blue and red dots which represents downregulated and upregulated piRNAs respectively. Dark blue indicates piRNAs species that did not show significant change in expression.

**Table 2:** Differential piRNA expression. Expression of piRNAs identified using piRBase from the indicated samples was compared. Number of upregulated or downreulated piRNAs was calculated for the sample indicated by * in comparison to the second sample for each comparison.

| Samples Compared | No. of Upregulated piRNAs | No. of Downregulated piRNAs | No. of piRNAs showing no change | Total piRNAs in the two samples |
| --- | --- | --- | --- | --- |
| G1 Tubulin-Gal4 DNMT3L* vs UAS-DNMT3L | 2358458 | 2265540 | 4975253 | 9599251 |
| G5 Tubulin-Gal4-DNMT3L* vs UAS-DNMT3L (ovary) | 2260619 | 2039478 | 20587716 | 24887813 |
| G5 Tubulin-Gal4-DNMT3L* vs G1 Tubulin-Gal4DNMT3L | 842349 | 517084 | 4228395 | 5587828 |
| G5 Tubulin-Gal4-DNMT3L* vs UAS-DNMT3L (embryo) | 2246938 | 2180640 | 20460235 | 24887813 |

Using the piRBase database, differential expression (overexpressed shown in green and downregulated in red) of ovarian piRNAs in G1 and G5 Tubulin-Gal4-DNMT3L as compared to control UAS-DNMT3L control transgenic flies was examined by plotting Volcano (Figure 3B, C, D) and Circos plots (Figure 4). Various piRNAs mapping to multiple loci on all Drosophila chromosomes were found to be misregulated in Tubulin-Gal4-DNMT3L flies of both G1 and G5 generation. We noticed a higher frequency of differential expressed piRNAs mapped to chromosomal ends. Interestingly, several overexpressed and downregulated piRNAs mapped to the same regions at several loci across the four Drosophila chromosomes. To further investigate the observation that both upregulated and downregulated piRNAs species map to same genetic regions at several loci (Figure 4), we decided to examine mapping of differentially expressed piRNAs at specific genetic loci. Based on the data of differential expressed genes in transgenic Tubulin-Gal4-DNMT3L flies [18], we selected a few upregulated (*Piwi*, *Vasa*, *Kdm4a*) and downregulated (*Abd-A*, *Wnt2*) genes for further analysis. As can be seen in Figure 5A, amongst the piRNAs expressed from the Piwi locus in the ovarian tissue, all piRNAs mapping to the intronic regions were upregulated in G5 transgenic Tubulin-Gal4-DNMT3L flies, whereas all the piRNAs mapping to the exonic regions of Piwi were downregulated as compared to the control UAS-DNMT3L flies. This was also observed for the piRNAs misregulated in the G5 transgenic Tubulin-Gal4-DNMT3L Drosophila embryos (Supplementary Figure S4).

**Figure 4:**
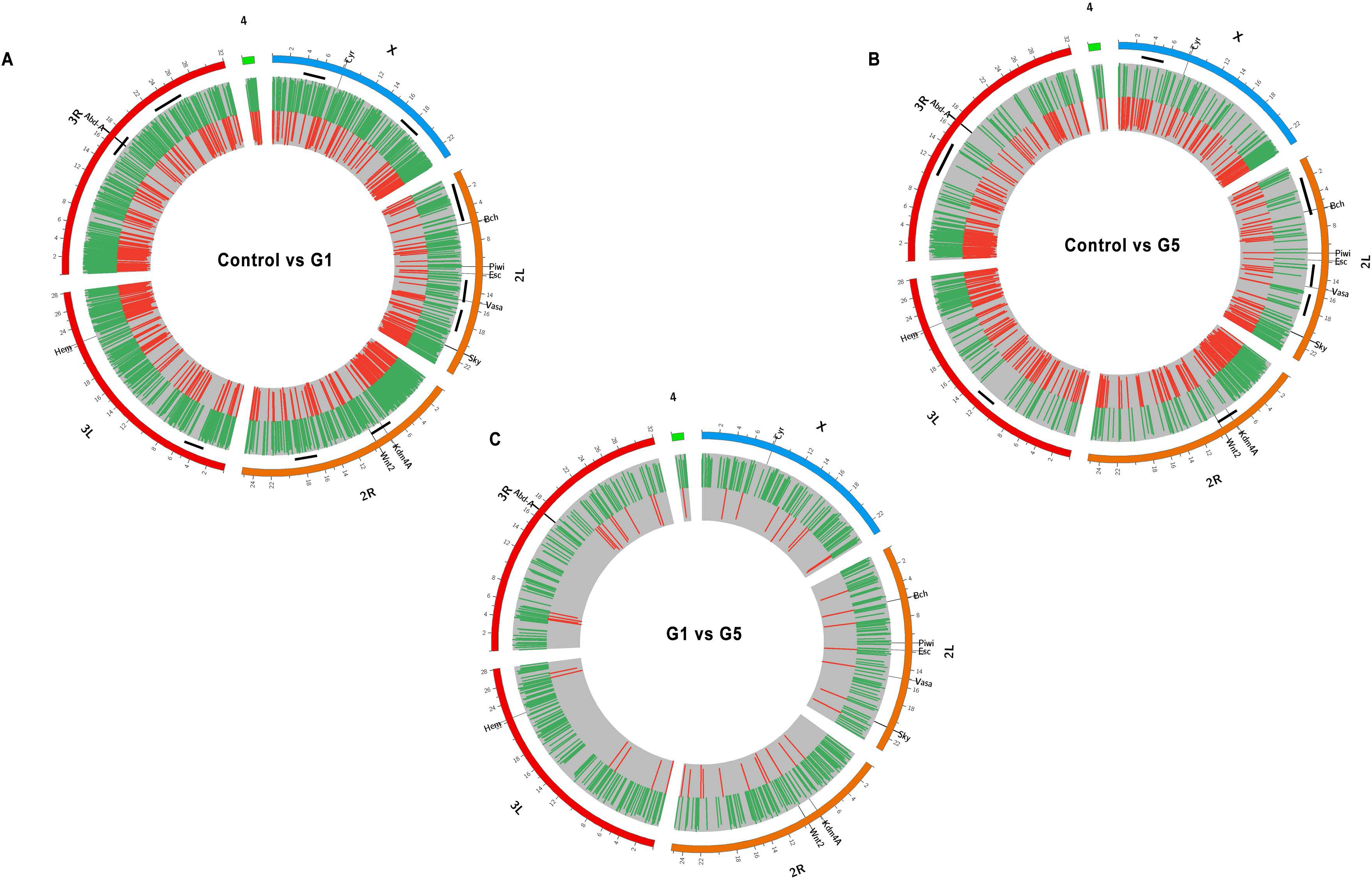
Circos plot mapping of differentially expressed piRNA in transgenic DNMT3L Drosophila. Differentially expressed piRNAs from ovarian tissue was mapped to different Drosophila chromosomes. A. Comparison of control UAS-DNMT3L vs G1 Tubulin-Gal4-DNMT3L; B. Comparison of control UAS-DNMT3L vs G5 Tubulin-Gal4-DNMT3L; and C. G1 vs G5 Tubulin-Gal4-DNMT3L. The expression values are in Log2 scale. Green and red lines represent overexpressed and downregulated piRNAs respectively in G1 (A.), G5 (B. & C.) flies. piRNAs expression from within specific genetic loci are shown in the outermost track. Black lines below ideogram in A & B represents differentially expressed piRNA.

**Figure 5:**
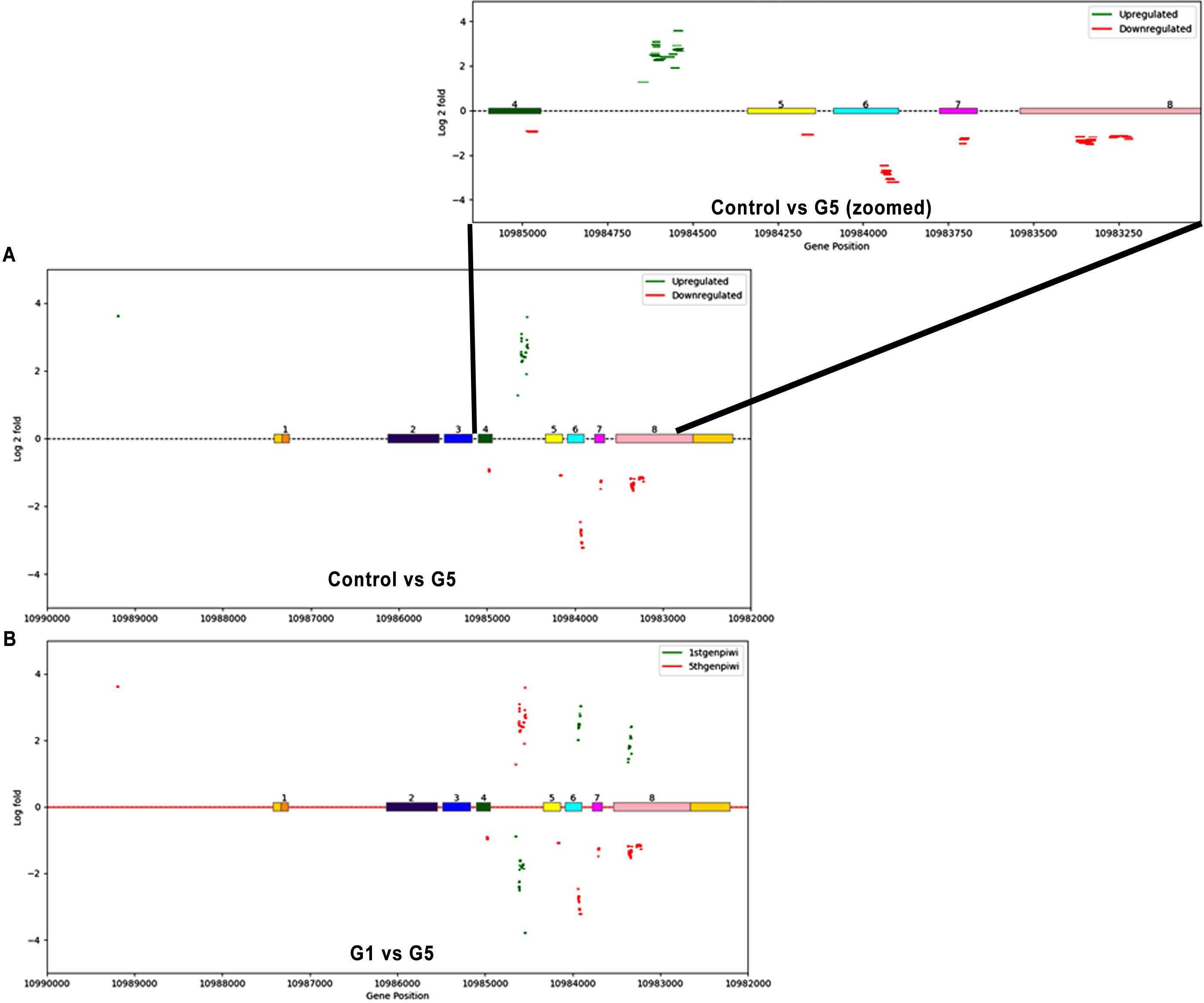
Representative images showing the mapping of differentially expressed piRNA clusters within the *Piwi* gene locus: A. piRNAs upregulated (green) or downregulated (red) in the ovarian tissue of G5 Tubulin-Gal4-DNMT3L transgenic flies as compared to control UAS-DNMT3L flies. Inset shows a zoomed representation of a region in A. B. piRNAs mapping to the same locus were upregulated (green) in G5 Tubulin-Gal4-DNMT3L but downregulated (red) in G1 Tubulin-Gal4-DNMT3L transgenic flies. The exonic regions of the Piwi gene are shown as coloured horizontal rectangles. Dashed line indicates intronic region. Number below X-axis denote the genomic position within the Drosophila genome. Differential expression is plotted for Log2fold values.

All the piRNAs mapping to the *Piwi* gene and differentially expressed in the ovarian tissue of G5 transgenic Tubulin-Gal4-DNMT3L Drosophila were also found to be differentially expressed in G1 transgenic Tubulin-Gal4-DNMT3L Drosophila ovaries. However, none of the upregulated piRNAs in the G5 transgenic Tubulin-Gal4-DNMT3L flies were upregulated in G1 flies. Same was the case with downregulated piRNAs. On the contrary, all the piRNA that were upregulated in G1 were found to be down regulated in G5 and vice versa all downregulated piRNAs were upregulated in G5 (Figure 5B). Similar profile was observed for piRNA produced from the upregulated gene *Vasa* and *Kdm4A* (Supplementary Figure S5).

*Abd-A* and *Wnt2* genes are downregulated in G5 transgenic Tubulin-Gal4-DNMT3L Drosophila. Very few differentially expressed piRNAs mapped to the *Abd-A* gene locus. These were located almost 20 kb upstream of the Abd-A TSS and were downregulated in G5 transgenic Tubulin-Gal4-DNMT3L flies (Supplementary Figure S6). As observed for upregulated genes, these few piRNAs associated with Abd-A locus were upregulated in G1 (Supplementary Figure S6). Similarly, for the other downregulated gene *Wnt2*, the differentially expressed piRNAs also mapped to the upstream region of these genes. For *Wnt2* locus, both up and downregulated piRNAs were detected in G5 transgenic Tubulin-Gal4-DNMT3Lflies. Expression profile for these piRNAs in G1 was also observed to be inverse of what observed in G5 transgenic Tubulin-Gal4-DNMT3L flies (Figure 6). Overall, the piRNA expression profile in G1 was almost always opposite to what was observed in G5.

**Figure 6:**
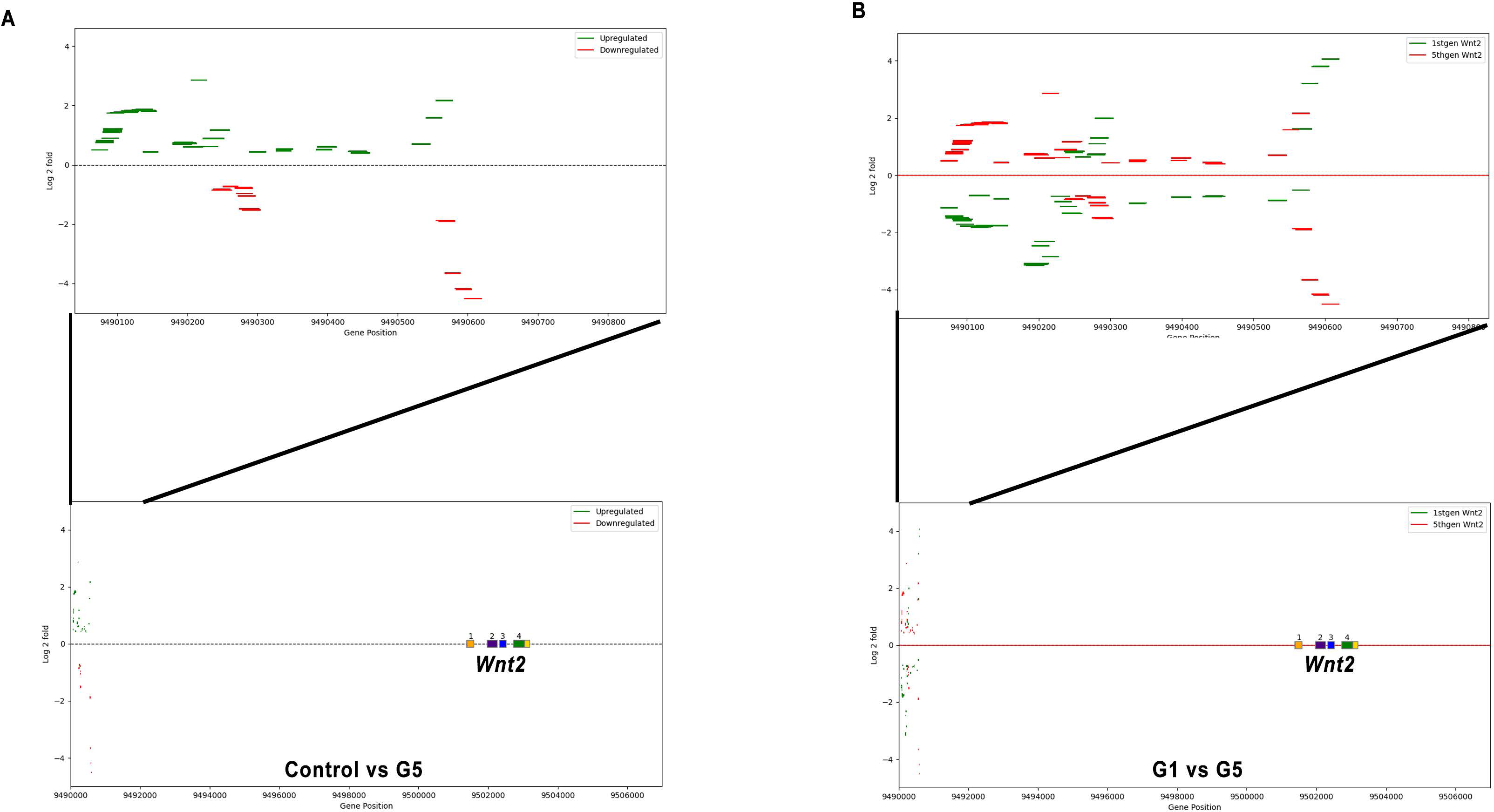
Representative images showing the mapping of differentially expressed piRNA clusters within the *Wnt2* gene locus: A. piRNAs upregulated (green) or downregulated (red) in the ovarian tissue of G5 Tubulin-Gal4-DNMT3L transgenic flies as compared to control UAS-DNMT3L flies. Inset shows a zoomed representation of a region in A. B. piRNAs mapping to the same locus were upregulated (green) in G5 Tubulin-Gal4-DNMT3L but downregulated (red) in G1 Tubulin-Gal4-DNMT3L transgenic flies. The exonic regions of the Piwi gene are shown as coloured horizontal rectangles. Dashed line indicates intronic region. Number below X-axis denote the genomic position within the Drosophila genome. Differential expression is plotted for Log2fold values.

## Discussion

piRNAs and the Piwi group of proteins have been identified as key components of epigenetic inheritance [27, 28, 29]. In the present study, we show that DNMT3L expressing transgenic Drosophila show progressive accumulation of epimutations [18], the population of piRNAs change dynamically across generations. In the light of our finding that DNMT3L interacts with Piwi protein and its collaborator Histone H1, a histone involved in higher order chromatin organisation, we propose that inheritance of non-genetic information results from multiple layers of epigenetic circuits.

### Interaction of DNMT3L with components of higher order chromatin organisation

The epigenetic modulator protein DNMT3L is known to function through its interaction with other *de novo* methyltransferases and core nucleosomal protein, Histone H3. Here, we show that DNMT3L also interacts with histone H1. Histone H1, unlike the core histones, has been correlated with higher order chromatin organisation including chromatin loops and chromosomal domains [30, 31, 32, 33]. Higher order chromatin organisation has been shown to be involved in synchronised expression of multiple genes, bringing regulatory elements like enhancers closer to gene promoter, etc. Moreover, modulation of gene expression by higher order chromatin organisation is cell-specific and depends upon environmental cues [34, 35]. Our finding that DNMT3L interacts with Histone H1 in addition to histone H3 indicates that DNMT3L has functions in addition to its role in modulating chromatin organisation through core histone modifications. It is possible that DNMT3L is also responsible for modulation of epigenetic modifications on histone H1. Further work would also be required to examine whether the interaction of DNMT3L with histone H1 and histone H3 regulate the same or different set of genes.

### Collaboration of piRNAs, Piwi and DNMT3L in epigenetic inheritance

Ectopic expression of DNMT3L causes the accumulation of epimutations across multiple generations and leads to development of melanotic tumors in 5^th^ generation of Tubulin-Gal4-DNMT3L transgenic flies. This accumulation of epimutations was dependent on the function of Piwi protein as the progeny of transgenic Tubulin-Gal4-DNMT3L crossed with Piwi mutant flies did not show melanotic tumors [18]. Moreover, expression of Piwi protein was highly upregulated in melanotic tumor containing G5 transgenic Tubulin-Gal4-DNMT3L 3^rd^ instar larvae. Here, we show that DNMT3L protein directly interacts with the Piwi protein. This could suggest that DNMT3L might be regulating the function of Piwi at multiple levels.

Piwi group of proteins have been shown to be predominantly involved in germline development and formation of gametes in several animal species [36, 37]. These proteins are known to perform their function through their association with piRNAs, [38, 39]. piRNAs were initially identified as transcriptional regulators of transposable elements but have subsequently been shown to target protein coding mRNAs and lncRNAs [40, 41]. While piRNAs are predominantly present in germ cells they are also synthesized in other cell types and their profile is considered as component of the epigenetic signature of a specific cell [42, 43, 44]. In this study, we show (i) piRNA profile in each generation was unique; (ii) piRNA profile of transgenic Tubulin-Gal4-DNMT3L flies in different generations was different from control UAS-DNMT3L flies; and (iii) piRNA profile in G1 and G5 transgenic Tubulin-Gal4-DNMT3L flies was almost a mirror-image of each other. These changes in piRNAs profile of transgenic Tubulin-Gal4-DNMT3L flies in different generations parallel epigenetic changes in the flies leading up to the melanotic tumors in G5 3^rd^ instar larvae. piRNAs have been shown to be intergenerationally inherited. Taken together with our previous observation that transcriptional reprogramming in transgenic Tubulin-Gal4-DNMT3L flies was progressive [18], we believe that DNMT3L through its interaction with Piwi modulates the profile of piRNAs, some of which are inherited to the transgenic Tubulin-Gal4-DNMT3L progeny in the next generation. piRNAs like other small RNA species are believed to play their regulatory role by bringing relevant epigenetic modifications to specific loci [9, 45]. Therefore, the possibility exists that this changed piRNA profile causes changes in epigenetic modifications (epimutations) in the subsequent generation, culminating into extreme loss of active histone marks (H3K4me2, me3 and H3K36me3) and causing melanotic tumors and subsequent death of these larvae in G5 [18]. However, it is still not clear as to why only 5-8% of the G5 Tubulin-Gal4-DNMT3L larvae showed melanotic tumors even though all others larvae also show loss of active histone modifications. It would also be interesting to understand why epimutations in Tubulin-Gal4-DNMT3L were only the active chromatin marks. It is possible that the cooperation of DNMT3L, Piwi, piRNAs and histone H1 works in conjunction of proteins like MLL and WDR5 involved in maintaining active chromatin states. Further work would be required to dissect out these interactions.

Finally, in this study we have uncovered a collaboration of an epigenetic modulator, DNMT3L, with a regulator of piRNAs, Piwi for inheritance of non-genetic information including piRNAs and active histone marks. Further studies to dissect out the mechanism based on this collaboration would help in unravelling the basis of non-genetic inheritance.

## Materials and Methods

### Fly stocks and crosses

Crosses to get transgenic UAS-DNMT3L and Tubulin-Gal4-DNMT3L Drosophila from various generations were performed as described in Basu et al 2016 [18].

### Pull-down and Immunoprecipitation assay

Pull-down and immunoprecipitation assays were performed by incubation in NETN buffer (20 mM Tris PH 8.0, 100 mM NaCl, 1 mM EDTA, 0.5% NP40 & protease inhibitors) followed by washing, elution of bound proteins by boiling in SDS sample buffer and analysis by western blotting For DNMT3L-Drosophila Piwi pull down, *E.coli* purified His-tagged DNMT3L (cloned in pET 28 a+) pre-equilibrated with Talon beads was used. For DNMT3L-histone H1 pull down, *E.coli* purified GST-Histone-H1 (Drosophila or human) or control GST bound to glutathione Sepharose beads (Amersham) were incubated with *E.coli* purified His-tagged Flag DNMT3L. Vice-versa, *E.coli* purified GST-tagged DNMT3L or control GST bound to glutathione Sepharose beads (Amersham) were incubated with *E.coli* purified His-Myc-Histone H1 (Drosophila or human protein). For pulldown assay with proteins purified from HEK293 cells (a kind gift from Dr Gayatri Ramakrishna, who obtained it from the Cell Culture Stock Centre at National Centre for Cell Science (NCCS), Pune), HA-DNMT3L and SFB-Piwi constructs were used. For immunoprecipitation assay, Drosophila ovaries lysate was incubated with anti-DNMT3L antibody or anti-Piwi antibody and the western blot was probed with an appropriate antibody.

### Source of antibodies

Following antibodies were used in the present study: anti-Piwi (sc-390946, Santa Cruz Biotechnology), anti-Flag (F1804, Sigma), anti-DNMT3L (ab194094, Abcam), anti-Rabbit IgG (ab171870, Abcam), anti-Mouse IgG (10400C, Sigma), anti-HA (ab9110, Abcam), anti-GST (ab19256, Abcam), anti-MYC (ab9106, Abcam).

### sRNA purification

Small RNAs were purified from ovaries and embryos using miRNeasy mini kit (cat no 217004, Qiagen) and RNeasy MinElute clean-up kit (cat no 74204, Qiagen). Ovaries and embryos were homogenized in Qiazol reagent as per manufacturer’s instructions. RNA was eluted in RNase-free water. The purification quality of small RNA was examined on agarose gel.

### Library preparation and sequencing

Illumina Trueseq Small RNA Library Prep kit was used to prepare the libraries for small RNA sequencing. For library preparation, 2 µg of small RNA (quantified by Nanodrop and Qubit) was used as starting material for all the samples. In modification to the Trueseq protocol, as 2srRNAs forms ∼90% of reads, its PCR amplification was blocked by terminator oligo block after 3’ adapter ligation [46]. Prepared libraries were selected for 140 to 160bp products by PAGE and purified. Final purified libraries were checked for quality and pooled in equimolar concentration and loaded on Illumina Nextseq 2000 machine for sequencing. Sequencing was performed in paired-end manner with 50bp coverage. Approximately 30 million reads for each sample were achieved.

### Bioinformatic analysis

The quality of reads was assessed by Fast QC. Adapter and low-quality reads and bases were removed using Trim Galore and BBDuk. Reads matching to 2srRNA removed by using bowtie and Samtools. Differential expression was performed using DESeq2 after quantification of reads. For checking the composition of various small RNAs, Unitas software was utilized. Circos plots were made using circos-0.69-9 version [47]. For plotting various piRNAs mapping on the genomic locus of different genes, python code was created (utilises log2fold, genomic coordinates columns in the excel sheet). Code used for plotting Piwi gene is provided in the supplementary files.

## Acknowledgements

We acknowledge Dr. Jamy C. Peng for providing vector containing Flag-Piwi for our experiments. We acknowledge Dr. Rohit Joshi and Dr. N. G. Prasad for use of their Drosophila facilities. We thank Dr. Shreekant Verma for assisting with fly experiments. We thank Sachin Bhakt for helping out in developing piRNA mapping tool for particular gene locus in the study. R.R is the recipient of Junior and Senior Research Fellowships of the Department of Biotechnology and was registered with the Manipal University for his PhD degree. SK is a JC Bose Fellow of the Department of Science and Technology, India and supported by grants from CSIR and DBT, Government of India.

