## Supplementary figures for "DNMT3L interacts with Piwi and modulates the expression of piRNAs"

### Supplementary figure S1

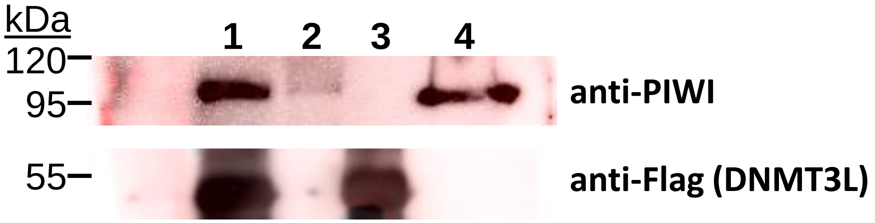


**Supplementary Figure S1: DNMT3L protein interacts with Piwi.** Western blot analysis was performed for DNMT3L-associated proteins by incubating recombinant 6X His-Flag-DNMT3L with $w^{1118}$ adult Drosophila cell lysate and affinity pulldown with Talon beads. The blot was probed with anti-Piwi antibody. Pull down with Talon beads alone was used as control. 1. Pulldown with Talon beads on Drosophila Lysate + 6X His-Flag-DNMT3L; 2- Pulldown with Talon beads on Drosophila lysate alone; 3- Input 6X His-Flag-DNMT3L; Lane 4- Input Drosophila lysate.

#### Supplementary Figure S2

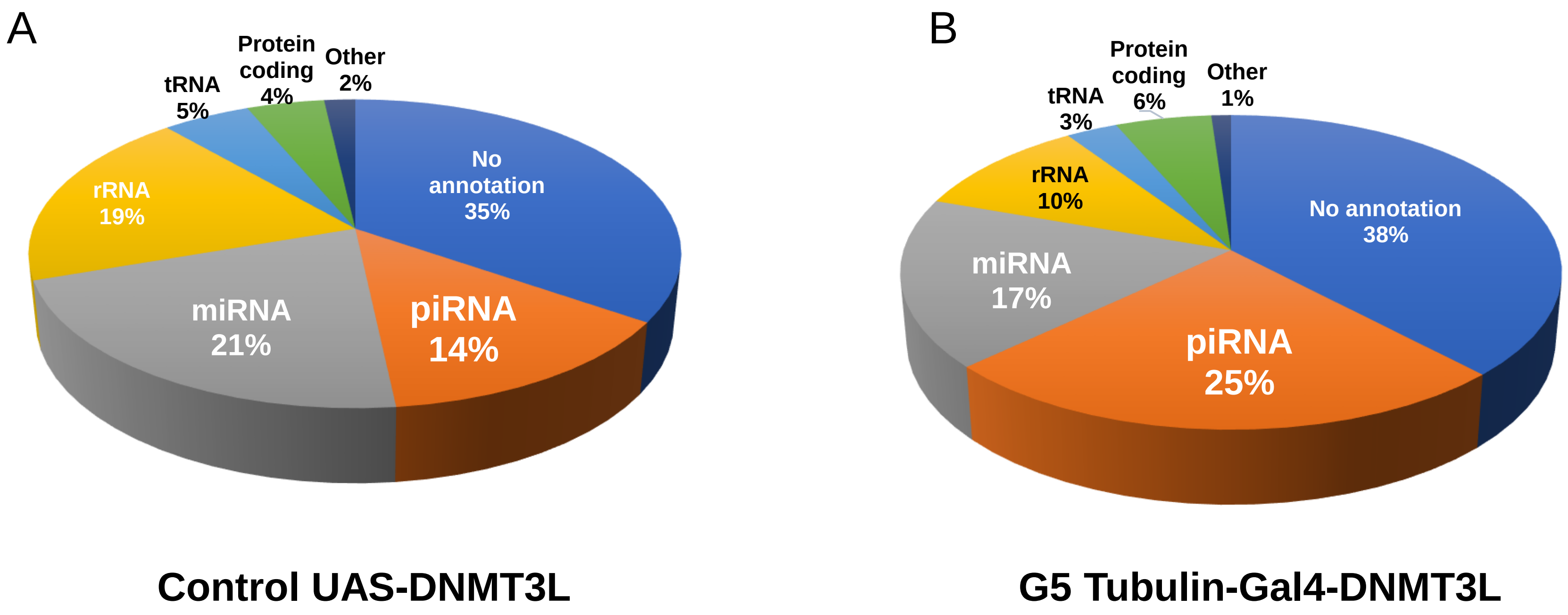


**Supplementary Figure S2: Composition of small RNAs (sRNA) in transgenic Drosophila embryo. A.** Pie charts representing the percent distribution of overall composition of small RNAs (sRNA) in the embryos isolated from **A.** UAS-DNMT3L control (without DNMT3L expression), and **B.** 5$^{th}$ generation of Tubulin-Gal4-DNMT3L (DNMT3L expressing) transgenic flies as identified in Unitas software.

### Supplementary Figure S3

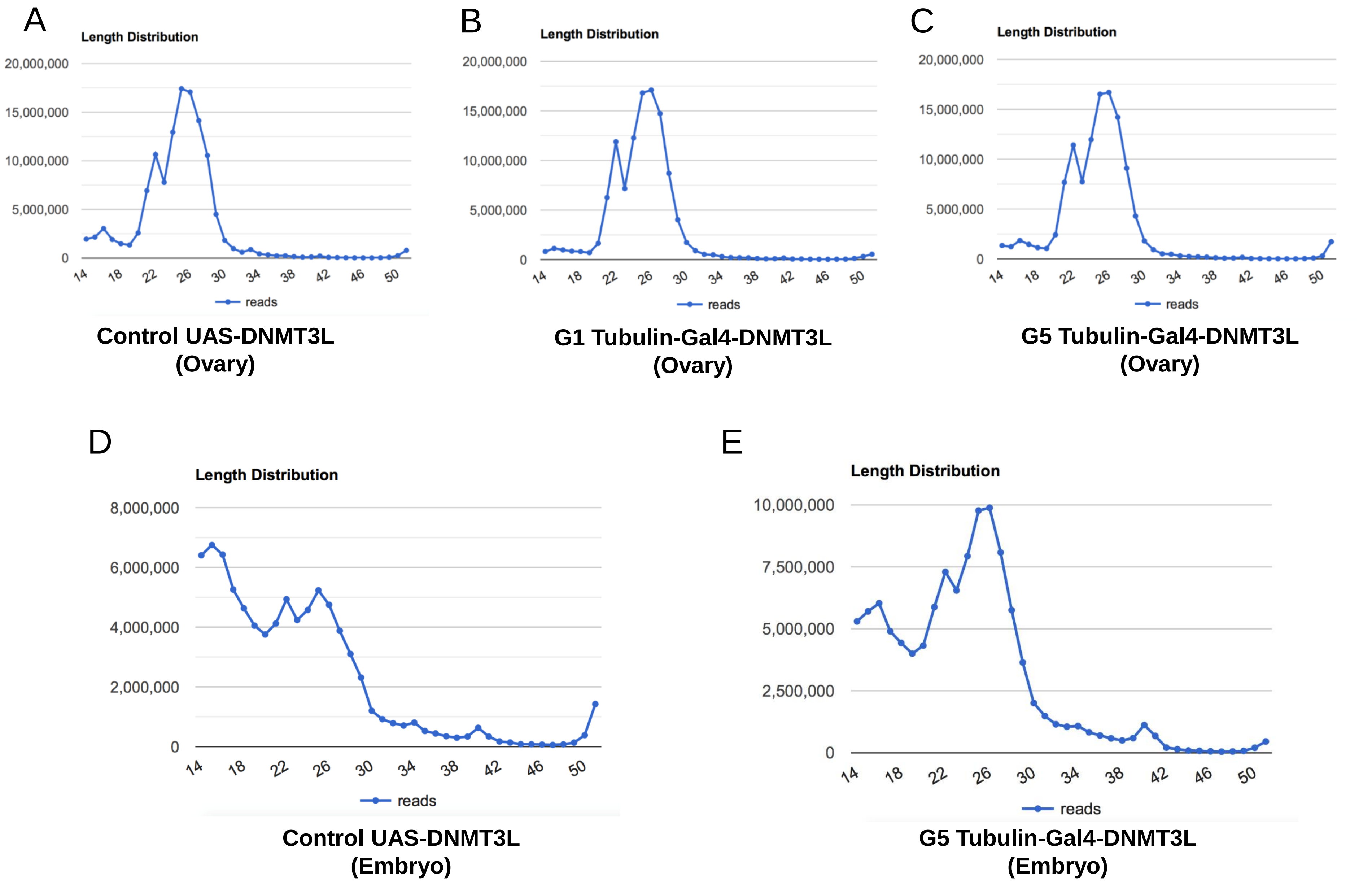


**Supplementary Figure S3:** The graphs illustrates the overall read lengths for all small RNA obtained in transgenic Drosophila as indicated below each panel.

**Supplementary Figure S4**

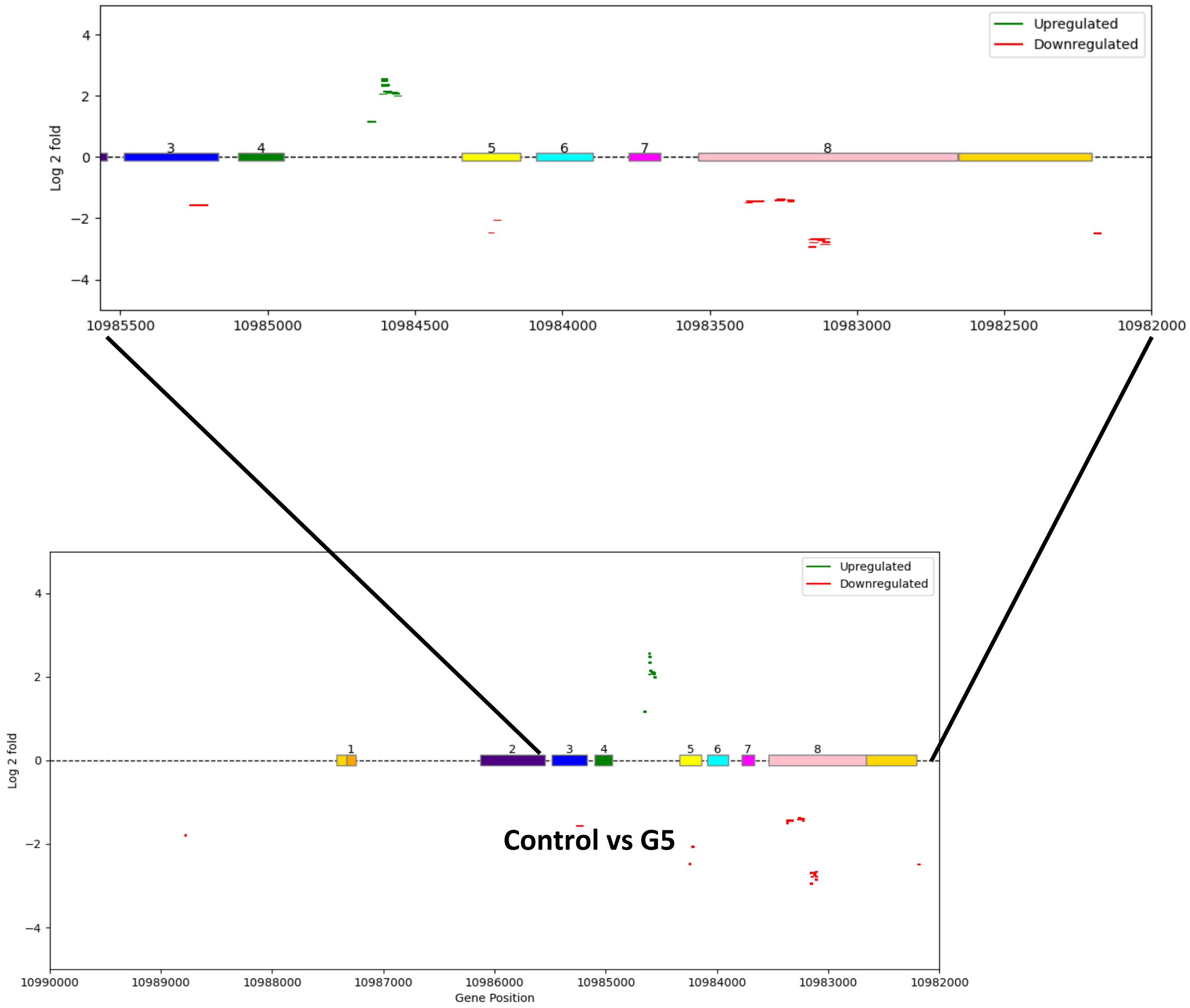


**Supplementary Figure S4: Representative images showing the mapping of differentially expressed piRNA clusters within the Piwi gene locus in embryonic tissue:** piRNAs upregulated (green) or downregulated (red) in the embryonic tissue of G5 Tubulin-Gal4-DNMT3L transgenic flies as compared to control UAS-DNMT3L flies. Inset shows a zoomed representation of a region. The exonic regions of the Piwi gene are shown as coloured horizontal rectangles. Dashed line indicates intronic region. Number below X-axis denote the genomic position within the Drosophila genome. Differential expression is plotted for Log2fold values.

### Supplementary Figure S5

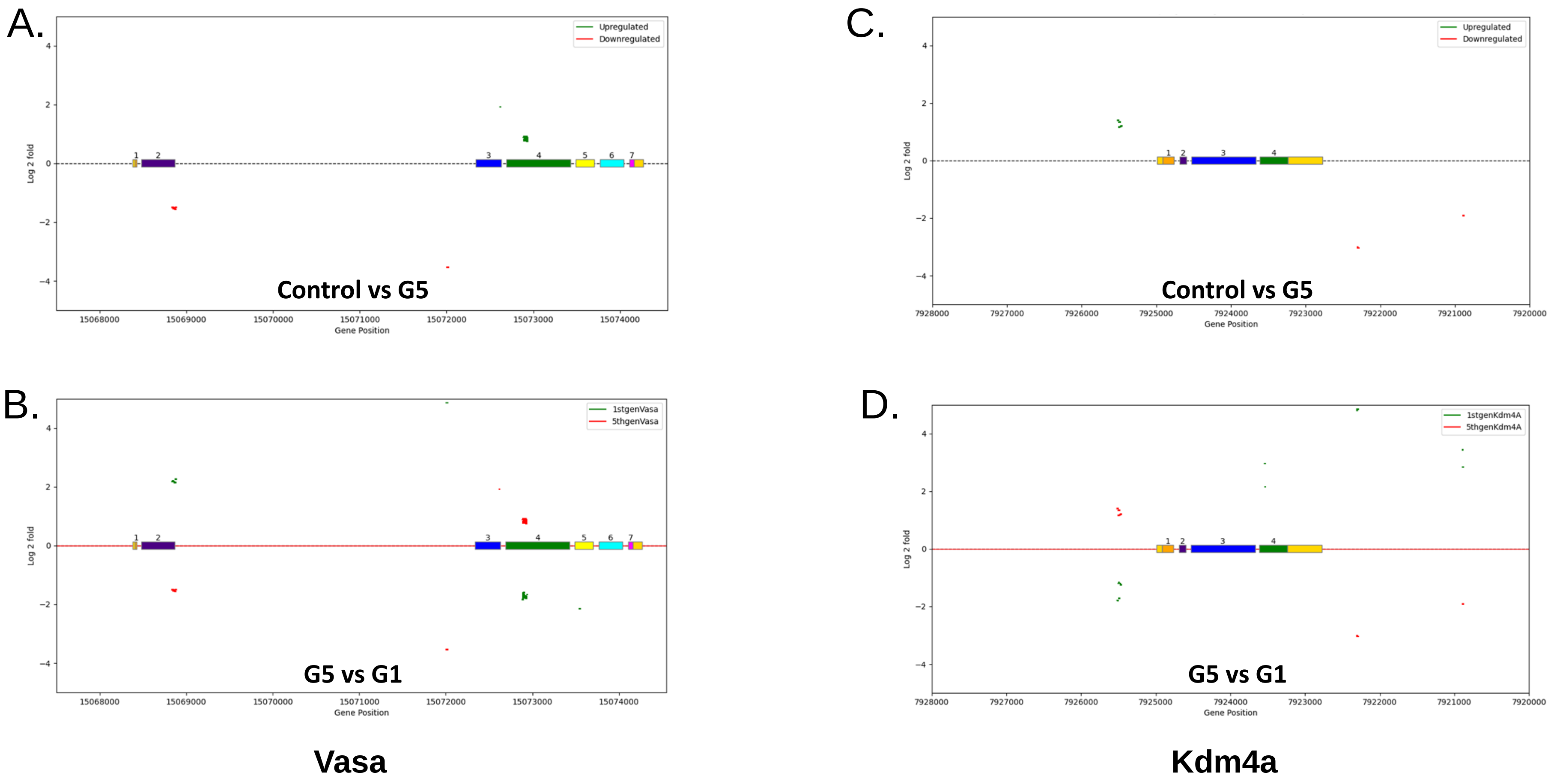


**Supplementary Figure S5: Representative images showing the mapping of differentially expressed piRNA clusters within the *Vasa* (A & B) and *Kdm4a* (C & D) gene loci :** piRNAs upregulated (green) or downregulated (red) in the ovarian tissue of: A. & C. G5 Tubulin-Gal4-DNMT3L transgenic flies as compared to control UAS-DNMT3L flies; B. & D. G5 Tubulin-Gal4-DNMT3L transgenic flies as compared to control G1 Tubulin-Gal4-DNMT3L flies. The exonic regions of the *Vasa* and *Kdm4a* gene are shown as coloured horizontal rectangles. Dashed line indicates intronic region. Number below X-axis denote the genomic position within the Drosophila genome. Differential expression is plotted for Log2fold values.

### Supplementary figure S6

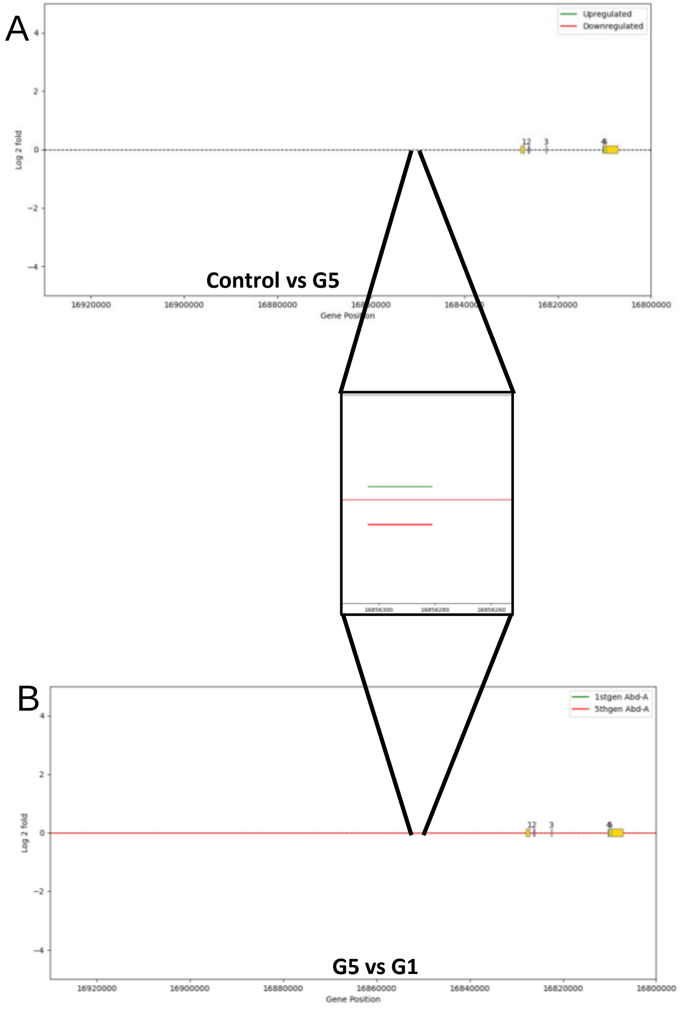


Abd-A

**Figure 6: Representative images showing the mapping of differentially expressed piRNA clusters within the *Abd-A* gene locus:** A. piRNAs upregulated (green) or downregulated (red) in the ovarian tissue of G5 Tubulin-Gal4-DNMT3L transgenic flies as compared to control UAS-DNMT3L flies. B. piRNAs mapping to the same locus were downregulated in G5 Tubulin-Gal4-DNMT3L but upregulated in G1 Tubulin-Gal4-DNMT3L transgenic flies (green represents G1 vs control and red represents G5 vs control). Inset shows a zoomed representation of a region in both A and B. The exonic regions of the *Abd-A* gene are shown as coloured horizontal rectangles. Dashed line indicates intronic region. Number below X-axis denote the genomic position within the Drosophila genome. Differential expression is plotted for Log2fold values.
